# The impact of rearing environment on *C. elegans*: Phenotypic, transcriptomic and intergenerational responses to 3D enriched habitats

**DOI:** 10.1101/2025.09.07.674770

**Authors:** Aurélie Guisnet, Nour Halaby, Maxime Rivest, Beatriz Romero Quineche, Michael Hendricks

**Affiliations:** McGill University; Independent Researcher; University of Ottawa

**Keywords:** *Caenorhabditis elegans*, Plasticity, Environmental enrichment, Rearing environment, RNA sequencing, Intergenerational effects

## Abstract

Environmental context profoundly influences organismal biology, yet laboratory studies often rely on simplified conditions that may not fully capture natural phenotypic repertoire. This exploratory study investigated how rearing environment affects various aspects of *Caenorhabditis elegans* biology by comparing worms cultured in three-dimensional decellularized fruit tissue scaffolds with those raised on standard two-dimensional agar plates. While fat content and feeding rate remained stable across conditions, other life history traits demonstrated varying degrees of plasticity in response to environmental context. We observed that scaffold-grown worms exhibited reduced body size, altered reproductive strategies, and mild enhancements in stress resistance, burrowing ability, swimming kinematics and exploratory behavior. RNA sequencing revealed distinct transcriptional profiles between scaffold-grown and agar-grown worms, with most changes arising within one generation. Some traits showed evidence of intergenerational inheritance. Our findings highlight the sensitivity of *C. elegans* biology to rearing conditions and underscore the importance of considering environmental context in interpreting laboratory results. This work sets the foundation for future research into the mechanisms underlying environmental adaptation and phenotypic plasticity in model organisms.

**Summary statement:** This study reveals how simple changes in environmental complexity can alter the development, behavior, and gene expression of laboratory animals.

## Introduction

Environmental conditions are fundamental drivers of biological development, behavior, and physiology across the animal kingdom. However, laboratory studies often rely on simplified, uniform conditions to facilitate control and reproducibility, potentially limiting our understanding of how environmental variability shapes biological processes and raising important questions about the breadth and applicability of our findings (Petersen et al., 2015; Würbel, 2000). This discrepancy is particularly evident in the study of model organisms like *Caenorhabditis elegans*, which are conventionally cultured on two-dimensional agar plates with uniform bacterial lawns—a stark contrast to their wild, dynamic, three-dimensional environments rich in structural and microbial diversity (Brenner, 1974; Schulenburg & Félix, 2017).

Previous research has underscored that minimalistic laboratory conditions can significantly affect gene expression patterns and behaviors, potentially obscuring the full spectrum of an organism’s biological responses (Alfred & Baldwin, 2015; Voelkl et al., 2020). Altering the rearing environment, even within controlled laboratory settings, can provide insights into the adaptability and plasticity of biological systems, shedding light on gene-environment interactions that are otherwise overlooked. In our prior work, we introduced a novel method for cultivating *C. elegans* within three-dimensional (3D) decellularized fruit tissue scaffolds, aiming to create a more complex, enriched living environment while maintaining experimental control (Guisnet et al., 2021a). This approach allowed us to observe behaviors and phenotypes that differed from those exhibited under standard laboratory conditions, suggesting that environmental context plays a significant role in shaping biological outcomes. Specifically, worms displayed a preference for the enriched 3D environment and demonstrated complex dauer behaviors not commonly observed in conventional settings. Despite these initial findings, the broader implications of raising *C. elegans* in alternative environments remain largely unexplored.

The present exploratory study aims to address these questions by examining the effects of the rearing environment on a suite of phenotypic and behavioral traits in *C. elegans*. By comparing worms cultured in our enriched scaffolds with those raised under standard laboratory conditions, we seek to explore how environmental context influences biological processes. Additionally, we investigate whether the observed phenotypic changes are heritable, contributing to the understanding of how environmental factors can lead to intergenerational phenotypic variations (Rechavi & Lev, 2017).

By providing an exploratory characterization of the influence of the rearing environment on *C. elegans* biology, this study aims to lay the groundwork for future research into the mechanisms underlying these effects. Understanding the full spectrum of *C. elegans* biological responses to different environmental contexts is crucial not only for interpreting laboratory findings more accurately but also for leveraging this model organism to study fundamental questions about gene-environment interactions, phenotypic plasticity, and the evolution of adaptive traits. This work highlights the importance of incorporating environmental context into the interpretation of biological research to achieve a more comprehensive understanding of life sciences.

## Results

To ensure that phenotypic effects were not confounded with potential toxicity of the fruit tissue preparation process, we analyzed the composition of the decellularized scaffold matrix for traces of the decellularizing agent, sodium dodecyl sulfate (SDS), using high resolution magic angle spinning nuclear magnetic resonance spectroscopy (HR-MAS). We found no traces of SDS in the scaffold (Fig. S1), indicating that animals are not affected by detergent residues. Peaks identified were consistent with components of plant cell walls or their breakdown products.

To disentangle the immediate effects of the growing environment from potential intergenerationally inherited traits, we compared four groups throughout: worms with an agar ancestry and raised on agar (agar:agar); worms with an agar ancestry, but raised on scaffold (agar:scaffold); worms with a scaffold ancestry, but raised on agar (scaffold:agar); and worms with a scaffold ancestry and raised on scaffold (scaffold:scaffold) (Fig. 1). Ancestry was established by raising the parents of all tested animals for at least 10 consecutive generations in their respective scaffold or conventional agar habitat. This design allowed us to assess both the direct impact of the habitat and any epigenetic influences transmitted across generations.

**Figure 1.**
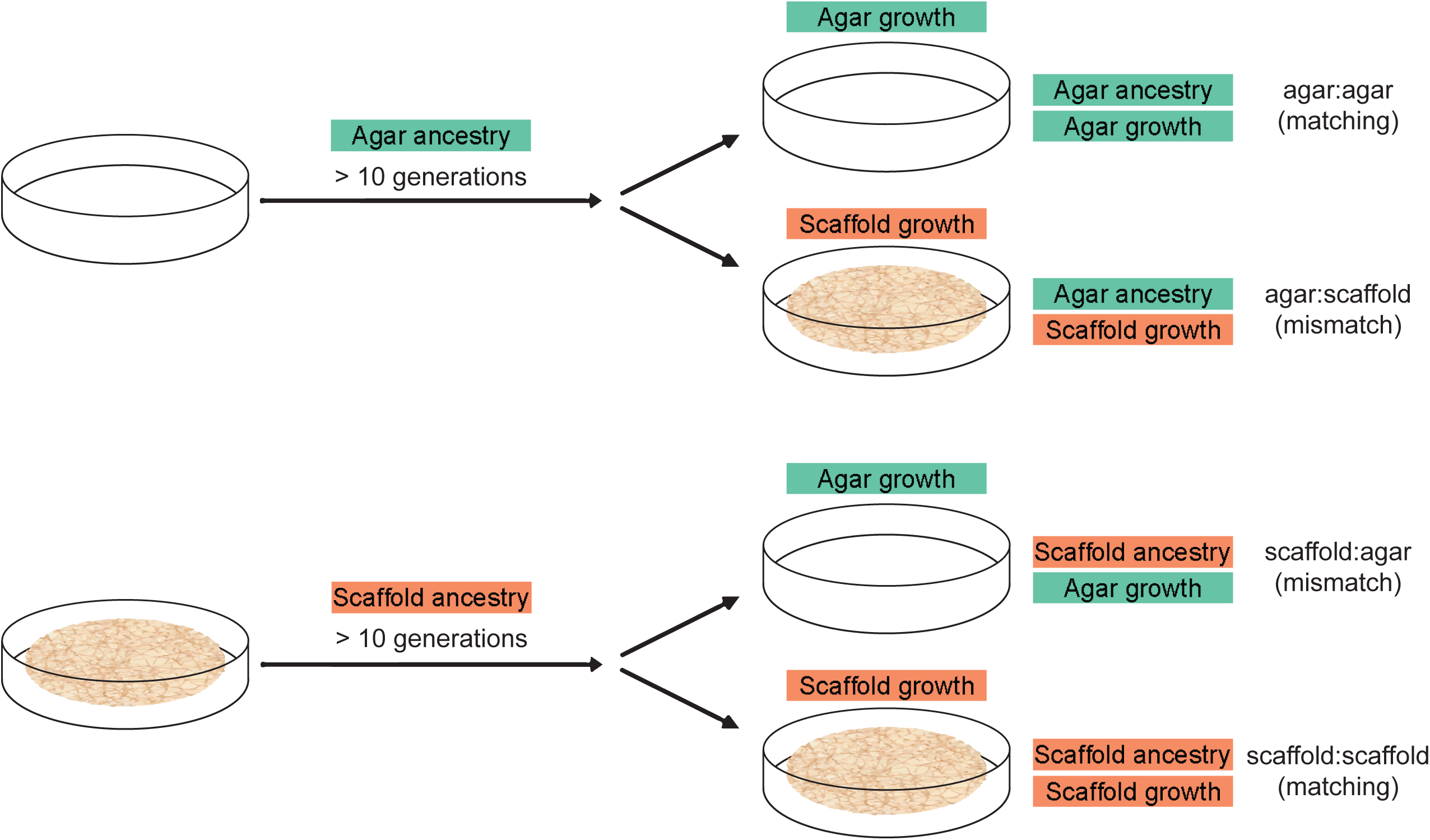
Experimental design investigating the effects of rearing environment and ancestry. Ancestry was established by raising the parents of all tested animals for at least 10 consecutive generations in their respective scaffold or conventional agar habitat. The growing habitat of experimental worms was established by moving gravid adults to either a matching or mismatched habitat to lay their eggs, leading to four experimental conditions.

### Pumping rate is not affected by habitat

We first investigated pharyngeal pumping rate, a key indicator of feeding behavior and overall health. The pumping rates across all conditions showed a relatively narrow range, with most values falling between 40 and 50 pumps per 10 seconds (Fig. 2A). This range is consistent with typical pumping rates reported for well-fed young adult *C. elegans* under standard laboratory conditions (Raizen et al., 1995).

**Figure 2.**
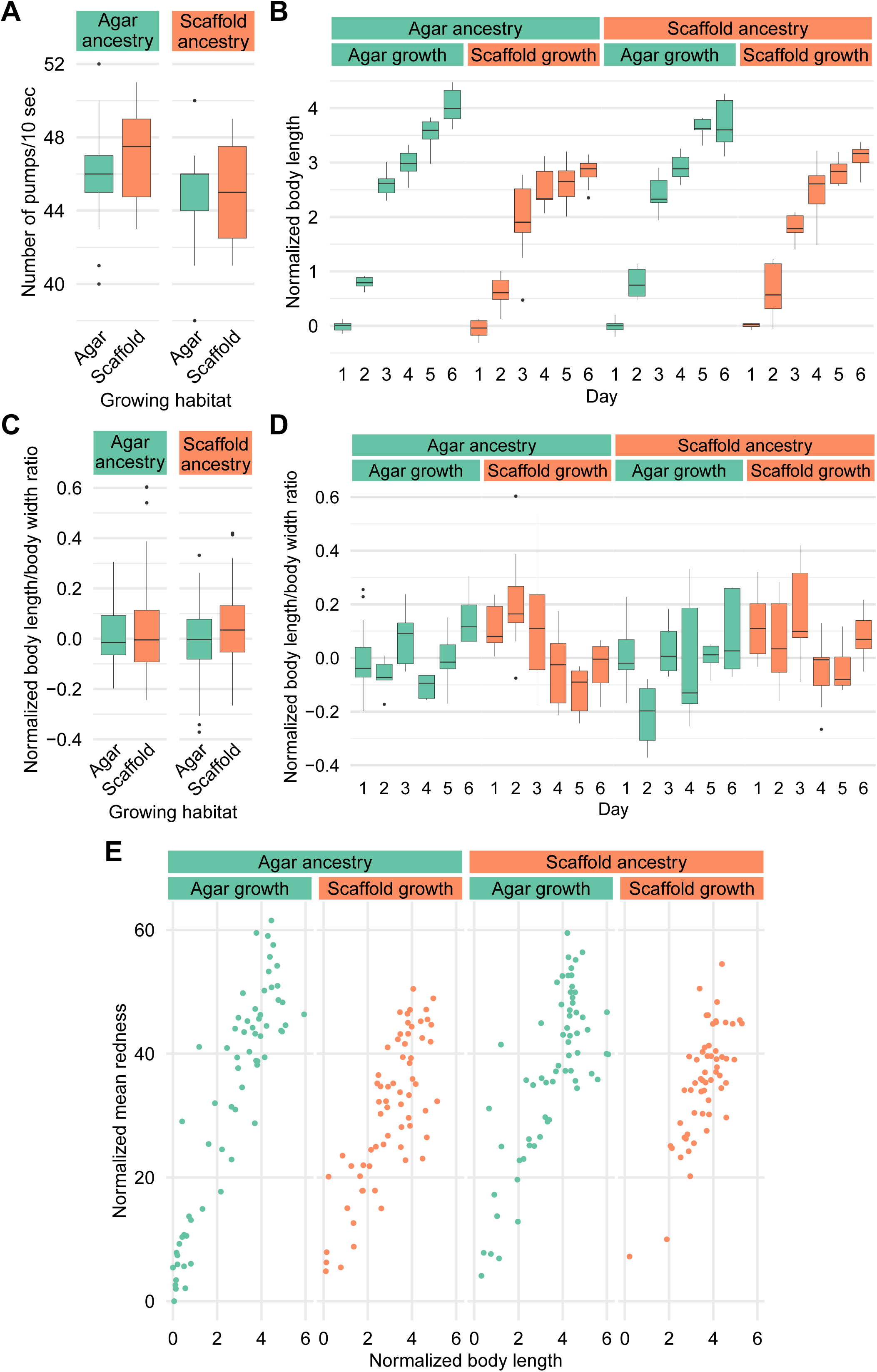
Environmental context modulates body size and developmental trajectories but not feeding rate or fat storage. (A) Number of pharyngeal pumps per 10 seconds. Sample size: agar:agar: n = 18; agar:scaffold: n = 16; scaffold:agar: n = 19; scaffold:scaffold: n = 19. (B) Body length by day. (C) Ratio of worm length by mean body width, all 6 days combined. (D) Ratio of worm length by mean body width by day. Sample size for (B-D): agar:agar: day 1: n = 13, day 2: n = 7, day 3: n = 12, day 4: n = 10, day 5: n = 11, day 6: n = 7; agar:scaffold: day 1: n = 8, day 2: n = 17, day 3: n = 10, day 4: n = 12, day 5: n = 8, day 6: n = 10; scaffold:agar: day 1: n = 12, day 2: n = 8, day 3: n = 11, day 4: n = 9, day 5: n = 6, day 6: n = 7; scaffold:scaffold: day 1: n = 7, day 2: n = 14, day 3: n = 11, day 4: n = 10, day 5: n = 6, day 6: n = 9. (E) Mean whole body redness by body length stained by Oil Red O. Sample size: agar:agar: n = 66; agar:scaffold: n = 65; scaffold:agar: n = 64; scaffold:scaffold: n = 57. All measured values in (B–E) are normalized to the mean of agar:agar worms on day 1.

### Developmental timing is altered by experience with the enriched environment

To characterize the effects on development, we imaged worms every 24 hours after egg laying over 6 days, and measured their morphological features.

Scaffold-grown worms, regardless of ancestry, showed reduced body length starting at day 3 compared to agar-grown worms (Fig. 2B). Mean body width revealed a similar pattern, with scaffold-grown worms generally being narrower than their agar-grown counterparts, although throughout the 6 days of development, and we observed for all groups a slight decrease in width after day 4 (Fig. S2A). Worm area and perimeter showed similar trends as body length (Figs S2C–D) and mid-body width similar to mean body width (Fig. S2B).

Body proportions of scaffold-grown and agar-grown worms remained similar on average (Fig. 2C), but scaffold-grown worms tended to be thinner in their early life and showed more variability in body proportions during early development. However, body proportions were more similar later in development (Figs 2D and S2E). Scaffold:agar worms had a strong tendency to be wider than agar:agar worms on day 2, but had similar proportions on other days (Fig. 2D).

### Fat content is not affected by habitat experience

To explore impacts on metabolism, we performed Oil Red O staining to assess major fat stores in worms of all life stages (O’Rourke et al., 2009).

We analyzed the staining intensity of individual worms relative to their body size. While the highest intensities were observed in agar-grown worms only, our results suggest that staining intensity was dependent on worm size across all conditions, where larger worms generally showed higher staining intensities (Fig. 2E). Importantly, no condition showed characteristics of calorically-restricted animals in major fat stores contents (Zhang & Mair, 2017).

### Environmental context and ancestry modulate total brood size and temporal patterns

We next tested whether the environment had an effect on total brood size and the distribution of egg-laying over time. We found that scaffold:scaffold worms had a smaller total brood than traditional agar:agar worms with a reduction mainly on the first and second days of egg-laying (Figs 3A and C). In addition, we found that both groups, once at the egg-laying stage, were unaffected by the egg-laying habitat (Fig. 3B).

**Figure 3.**
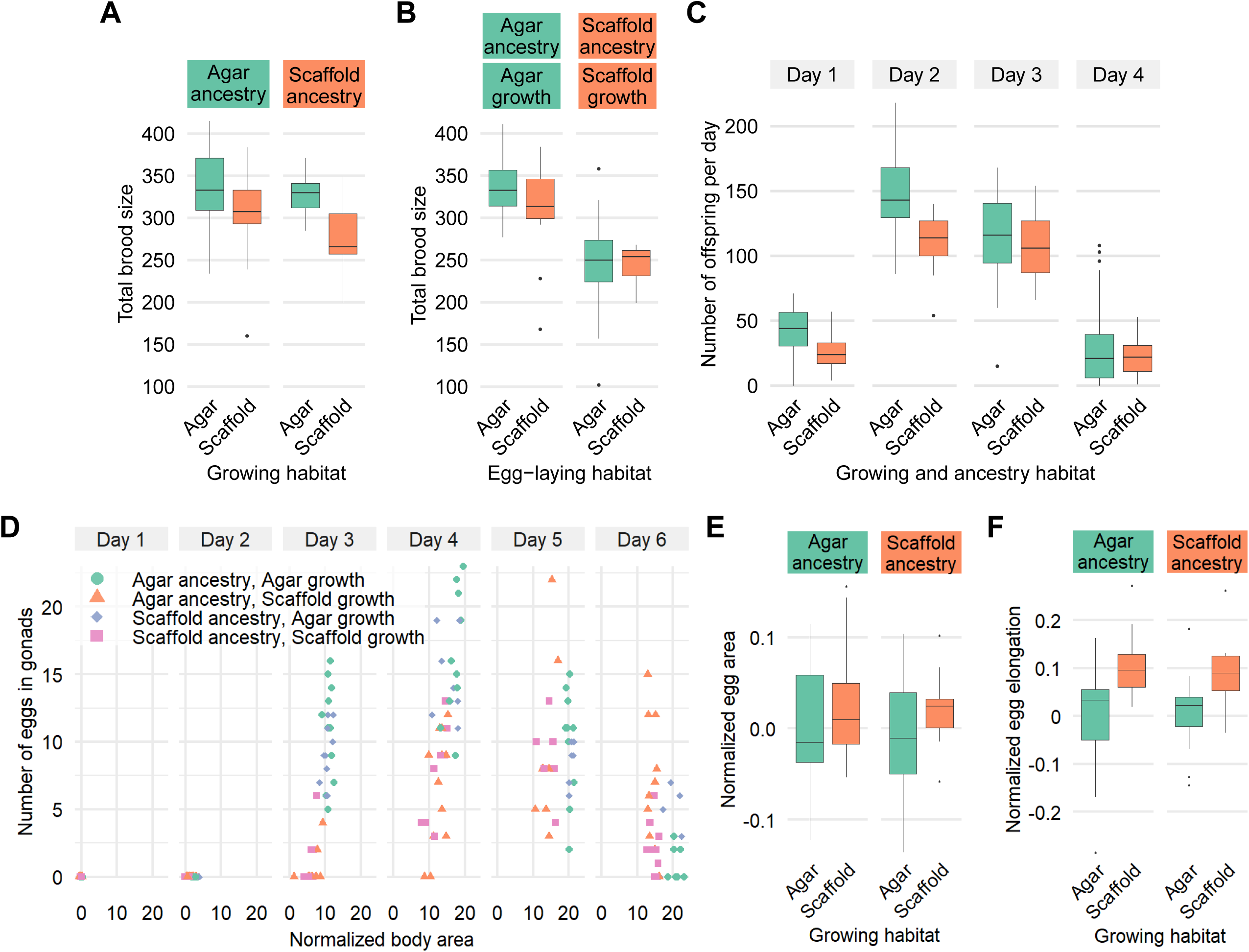
Rearing environment and ancestry influence reproductive strategies and egg morphology. (A) Total brood size. Sample size: agar:agar: n = 23; agar:scaffold: n = 26; scaffold:agar: n = 21; scaffold:scaffold: n = 13. (B) Total brood size for agar:agar and scaffold:scaffold worms, but once at the egg-laying stage, moved to a habitat matching or mismatched of their own. Sample size: agar on agar: n = 20; agar on scaffold: n = 14; scaffold on agar: n = 17; scaffold on scaffold: n = 8. (C) Number of offspring per day. Sample size for each day: agar: n = 43; scaffold: n = 21. (D) Number of eggs in gonads by worm body area normalized to the mean of agar:agar worms on day 1 by day. Sample size: agar:agar: day 1: n = 13, day 2: n = 7, day 3: n = 12, day 4: n = 10, day 5: n = 11, day 6: n = 7; agar:scaffold: day 1: n = 8, day 2: n = 17, day 3: n = 10, day 4: n = 12, day 5: n = 8, day 6: n = 10; scaffold:agar: day 1: n = 12, day 2: n = 8, day 3: n = 11, day 4: n = 9, day 5: n = 6, day 6: n = 7; scaffold:scaffold: day 1: n = 7, day 2: n = 14, day 3: n = 11, day 4: n = 10, day 5: n = 6, day 6: n = 9. (B) Egg area, and (C) egg elongation, normalized to the mean of agar:agar worms on day 1. Sample size (B–C): agar:agar: n = 14; agar:scaffold: n = 13; scaffold:scaffold: n = 12; scaffold:agar: n = 13.

For worms raised in a habitat mismatched to their ancestors, both agar:scaffold and scaffold:agar worms produced a similar number of offspring as agar:agar worms (Fig. 3A). This suggests a potential intergenerational effect of the agar habitat on reproduction, even when offspring are reared on enriched conditions.

### Egg production dynamics and morphology are affected the enriched habitat

To investigate further differences in development and brood size, we counted the number of eggs present in the gonads of individual worms over the 6 days after being laid. Eggs became visible in the gonads across all conditions only by day 3. The number of eggs present in the gonads were proportional to worm size, with worms accumulating more eggs in their gonads with size on day 3 and 4 (Fig. 3D). On days 5 and 6, when more size was more stable (Fig. 2B), the number of eggs in their gonads was also stable across all conditions, despite scaffold-grown worms having smaller body sizes (Fig. 3D).

While total egg area was similar across all conditions, eggs from scaffold-grown worms showed greater elongation, although variability was large across all groups (Fig.3F).

### Environmental context modulates burrowing behavior

To explore changes in neuromuscular performance and chemotaxis, we employed the burrowing assay described by Laranjeiro et al. (Laranjeiro et al., 2019) on young adult worms. This assay measures the ability of worms to navigate through a three-dimensional pluronic gel matrix towards an attractant (*E. coli* OP50, in our case).

All conditions showed a similar total fraction of worms to have reached the attractant (between 67.9–73.3%) after 3 hours, except for the agar:scaffold group which had only 50% of the animals reach the top (Fig. 4B). On the other hand, of the animals that reached the attractant, their median time to reach the top had a broad, overlapping range. Scaffold:scaffold worms had a median time to top of only 40 minutes, while the other groups took 85-90 minutes (Fig. 4A).

**Figure 4.**
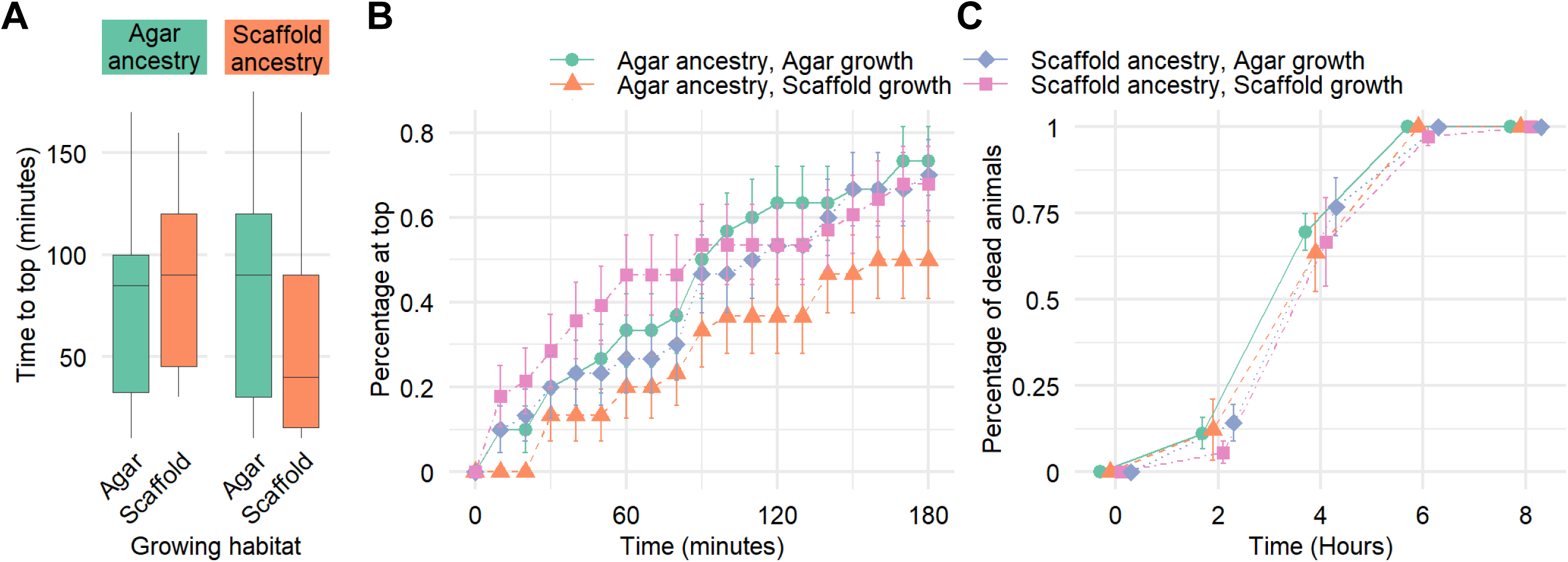
Burrowing ability and oxidative stress resistance are modestly modulated by environmental context and ancestry. (A) Time to reach the attractant at the top of the gel, successful animals only. (B) Total percentage of worms having reached the attractant located at the top of the gel for every 10 minute time-point over 180 minutes. Error bars represent standard error. Sample size (A–B): agar:agar: n = 30; agar:scaffold: n = 30; scaffold:scaffold: n = 28; scaffold:agar: n = 30. (C) Total percentage of dead worms for every 2 hour time-point over 8 hours. Error bars represent standard error. Sample size: agar:agar: n = 36; agar:scaffold: n = 35; scaffold:scaffold: n = 36; scaffold:agar: n = 36

Scaffold:scaffold worms particularly showed a larger percentage of worms reaching the top in the first 90 minutes (Fig. 4B).

### Scaffold-grown *C. elegans* exhibit a modest increase in oxidative stress resistance

To investigate the effects of the enriched habitat on stress resistance, we exposed young adult worms to oxidative stress using 3 mM H_2_O_2_ and monitored their survival every 2 hours over 8 hours. Our results suggest a slight enhancement in oxidative stress resistance for worms grown on scaffold. Particularly, only scaffold:scaffold animals were still alive at the 6-hour time point. No worm survived the entire 8-hour period (Fig. 4C).

### Swimming behavior is modulated by ancestry and experience

We further looked into behavioral differences by examining swimming behavior of young adults. We used methyl cellulose (MC) to create a range of viscous liquids: 0% (1 cP, same as water), 0.5% (5.6 cP, similar to milk) and 1% (20 cP, similar to vegetable oil). Higher viscosities were not tested as above 20 cP, worms exhibited crawl-like motion.

We started by looking into long-term swimming patterns by counting swimming bends in 5 seconds periods over 3 hours, as described in Ghosh & Emmons (2008). In the 0% MC solution, it was first evident that worms grown in a habitat mismatched from their ancestors spent more time being quiescent (0-2 bends per 5 seconds) (Figs 5A and S3A). However, of the periods where worms were swimming (5 bends or more per 5 seconds), scaffold-grown worms showed a slight increase in speed (Fig. 5C). There was no difference observed in the proportion of time spent in slow swimming (2-5 bends per 5 seconds) (Fig. 5B). A time series heatmap showed that worms grown in a habitat matching their ancestors particularly had no quiescent periods in the last 30 minutes of the 3 hour period (Fig. 5D). On the other hand, there was an observable increase in quiescence through time for animals grown in conditions mismatched from their ancestors (Fig. 5D). Still, all conditions showed slowing with time, although this trend was the strongest in scaffold:agar worms (Fig. S3B). In the more viscous solutions, worms would often be immobile at the side of the well, perhaps due to the increased surface tension. When considering only swimming periods, there was no marked difference in their speed in 0.5% MC, but a slight increase was observed in 1% MC again in scaffold-grown animals (Fig. 5C).

**Figure 5.**
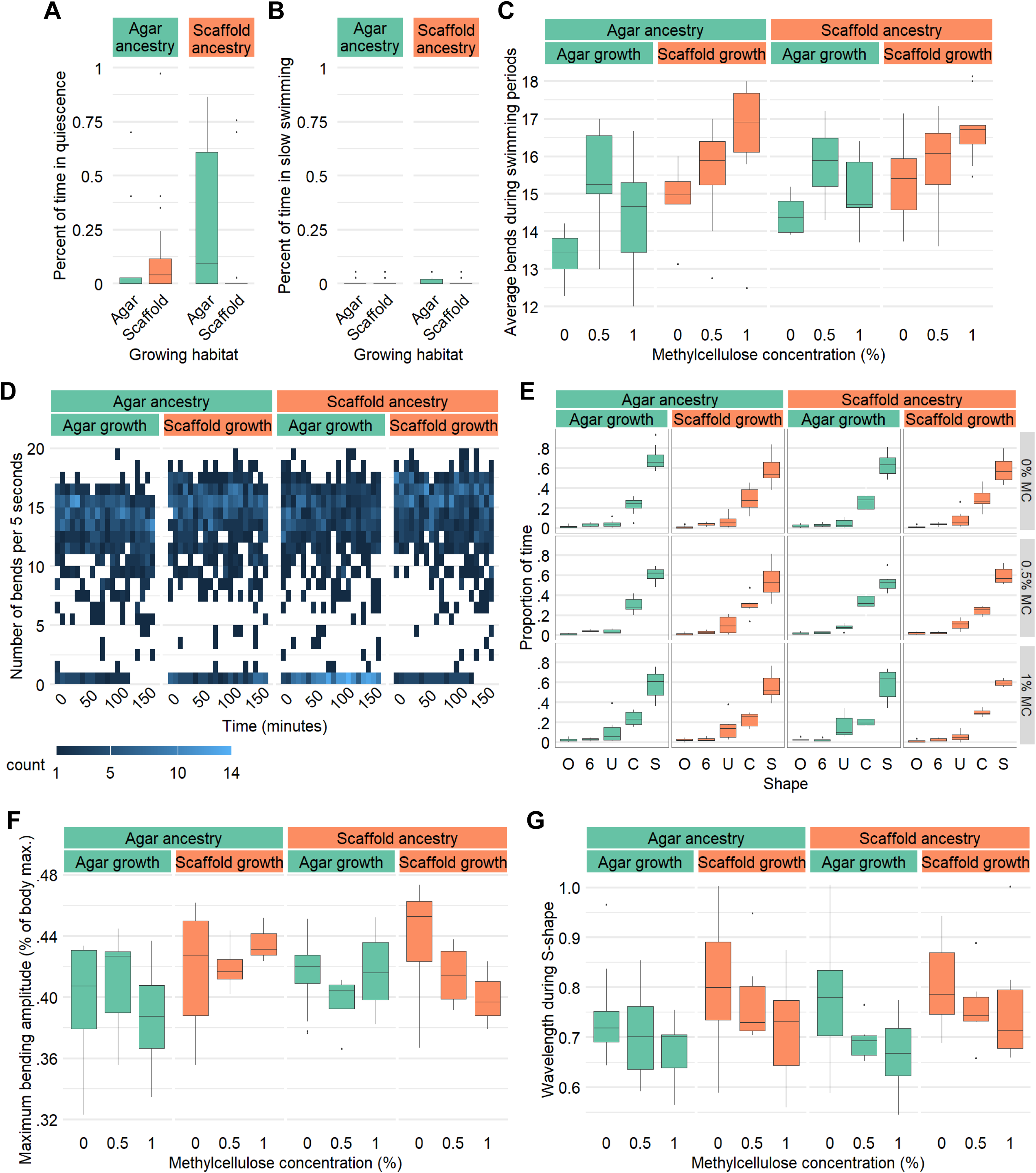
Swimming activity levels and kinematics are influenced by rearing environment and ancestry. (A) Percentage of the 3 hour experimental period spent in quiescence (≤2 bends/5 seconds). (B) Percentage of the 3 hour experimental period spent in slow swimming (3-4 bends/5 seconds). (C) Average number of bends in 5 seconds during swimming periods (≥5 bends/5 seconds) for different viscosities. (D) Heatmap of number of swimming bends per 5 seconds through time in 0% MC. Sample size (A–D): agar:agar: 0% MC: n = 17, 0.5% MC: n = 11, 1% MC: n = 9; agar:scaffold: 0% MC: n = 17, 0.5% MC: n = 9, 1% MC: n = 10; scaffold:agar: 0% MC: n = 17, 0.5% MC: n = 11, 1% MC: n = 10; s≥old:scaffold: 0% MC: n = 17, 0.5% MC: n = 7, 1% MC: n = 9. (E) Percentage of recorded frames where the worm body was in O, 6, U, C, or S-shape by viscosity. (F) Maximum bending amplitude in proportion to body length by viscosity (0 being perfectly flat and 1 being perfectly folded in half). (G) Number of wavelengths per body length for frames in S-shape by viscosity. Sample size (E–G): agar:agar: 0% MC: n = 15, 0.5% MC: n = 6, 1% MC: n = 6; agar:scaffold: 0% MC: n = 14, 0.5% MC: n = 6, 1% MC: n = 6; scaffold:agar: 0% MC: n = 15, 0.5% MC: n = 6, 1% MC: n = 6; scaffold:scaffold: 0% MC: n = 14, 0.5% MC: n = 6, 1% MC: n = 6.

We next recorded young adult worms swimming in liquid droplets for 1 minute in 0% MC, but for 30 seconds in 0.5% and 1% MC as they would rapidly settle on the edges of the droplet. We did not observe any difference in the proportion of time spent in different body shapes (Fig. 5E). Since scaffold-grown worms were on average smaller as adults than agar-grown worms, we calculated their bend amplitude as a proportion of their body length. This comparison revealed no marked difference (Fig. 5F). For frames where animals were in S-shape, no difference in body wavelength was observed (Fig. 5G).

### Exploratory behavior is mildly affected by experience with the scaffold habitat

To investigate influences on locomotion on a 2D surface, we analyzed the crawling behavior of young adult worms for 1 minute under three different food availability conditions: absence of food (none), low food density, and high food density.

Across all conditions, worms spent the majority of their time moving forward and spent slightly more time stationary than moving backward (Fig. 6A). However, the agar:agar worms in low food density, spent less time moving forward and more time stationary, resulting in nearly equal time allocation between these two states (Fig. 6A). The average speed during both forward and backward movement was not different between groups or food densities (Fig. 6B).

**Figure 6.**
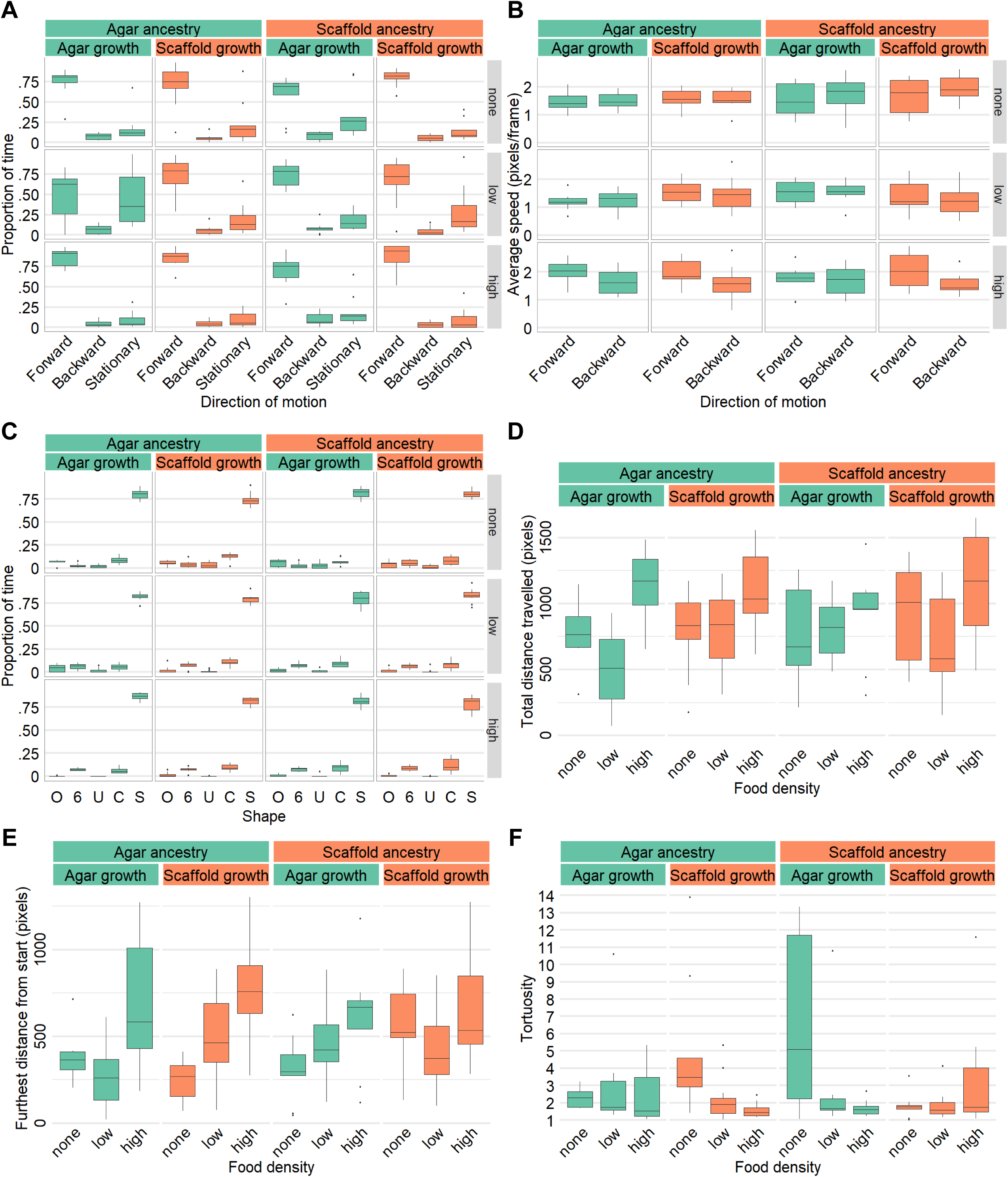
Crawling behavior is mildly affected by environmental context, ancestry, and food availability. (A) Proportion of time spent per recording in different directions of motion by food density. (B) Average speed in pixels per frame for forward and backward movement by food density. (C) Proportion of time per recording where the worm body was in O, 6, U, C, or S-shape by food density. (D) Total distance traveled in pixels by food density. (E) Furthest Euclidean distance reached from start point in pixels by food density. (F) Path tortuosity (total distance travelled by distance between start and end points) by food density. Sample size (A–F): agar:agar: no food: n = 8, low food density: n = 11, high food density: n = 10; agar:scaffold: no food: n = 9, low food density: n = 11, high food density: n = 10; scaffold:agar: no food: n = 9, low food density: n = 11, high food density: n = 9; scaffold:scaffold: no food: n = 9, low food density: n = 11, high food density: n = 10.

There was no difference in the proportion of time spent in different body shapes with S-shaped postures being overwhelmingly dominant and other shapes occurring infrequently and at similar low levels (Fig. 6C). This suggests no difference in the frequency of deep turns which are characterized by O, U and 6-shapes. Similarly, analysis of head bend movements revealed no differences in the frequency of bends (Fig. S4A) or their depth (Fig. S4B).

On average, the total distance travelled and the furthest distance reached did not differ considerably, except for scaffold:scaffold worms which travelled approximately 30% further in the absence of food (Figs 6D, E). While tortuosity was similar in the presence of food (low and high density), in the absence of food, worms raised in an habitat mismatched to their ancestors exhibited higher path tortuosity suggesting a prioritization of local search (Fig. 6F). In both low and high-density food environments, all groups reached their furthest exploratory point near the end of the recording period (Fig. S4C). In the absence of food, this trend remained true for scaffold:scaffold worms, but was slightly less pronounced for the other groups (Fig. S4C).

### Rearing environment drives mild, rapidly reversible changes transcriptional profiles

We investigated transcriptional changes by performing RNA sequencing on young adult worms. Principal component analysis revealed that rearing environment and ancestry were only mild drivers of gene expression differences, aligning with the subtle phenotypic changes observed in our study (Figs 7A–B).

**Figure 7.**
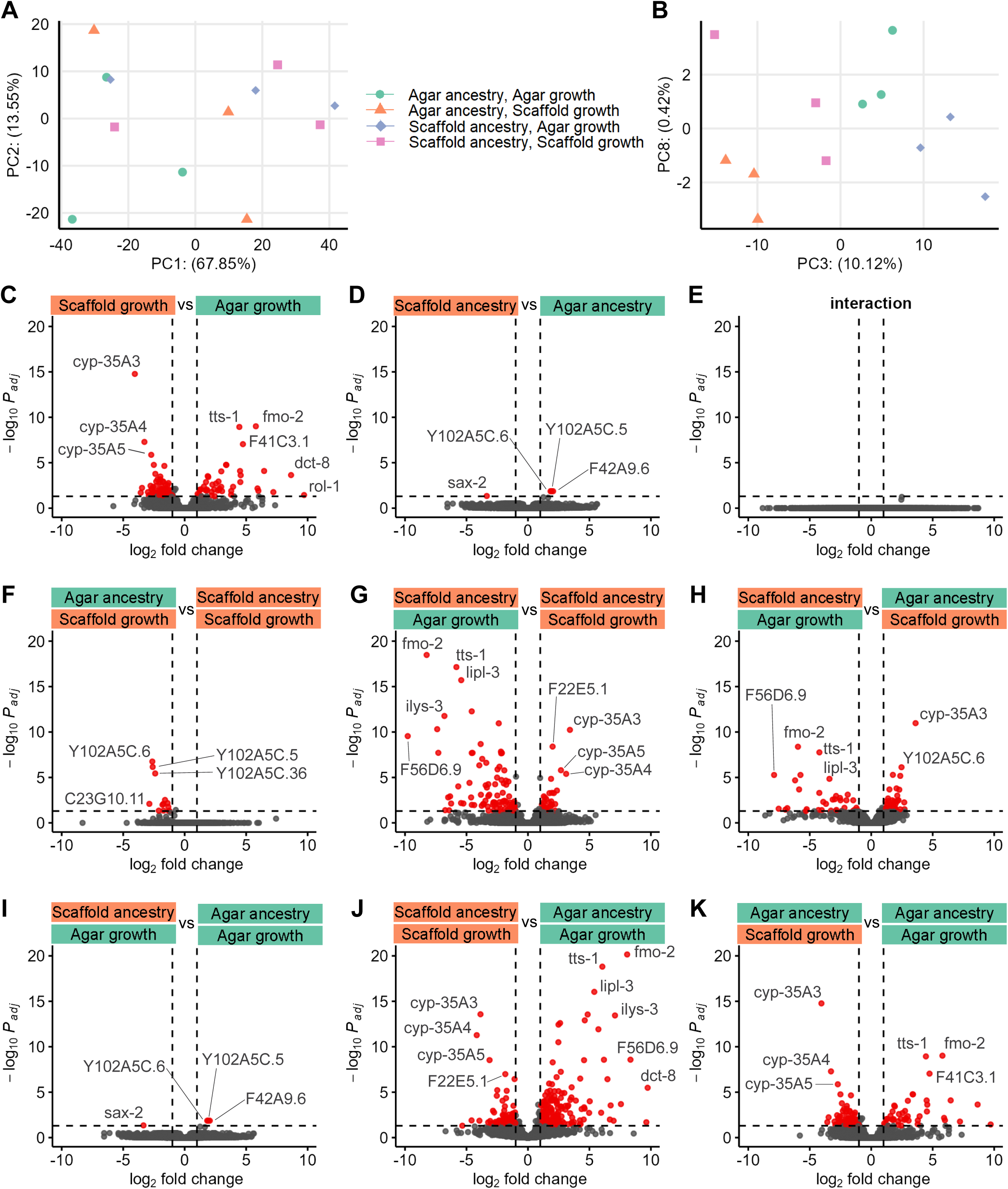
Transcriptional profiles reveal mild, rapidly reversible changes. Principal components (A) 1 and 2, and (B) 3 and 8, where each point represents a biological replicate of RNA sequencing. Volcano plots of factorwise differential expression analysis for (C) growth habitat, (D) ancestry habitat and (E) interaction term. Volcano plots of pairwise differential expression analysis for (F) agar:scaffold versus scaffold:scaffold, (G) scaffold:agar versus scaffold:scaffold, (H) scaffold:agar versus agar:scaffold, (I) scaffold:agar versus agar:agar, (J) scaffold:scaffold versus agar:agar and (K) agar:scaffold versus agar:agar groups. For (C–K), each dot represents a gene with red-colored being significantly differentially expressed between the two conditions by a log_2_fold change >|1| and a p-adjusted <0.05. Some genes of interest are labelled. 3 sequencing replicates per condition.

To explore the specific effects of the immediate growing environment and ancestry, we conducted factorial differential expression analysis. This analysis confirmed that the rearing habitat was the most significant factor influencing gene expression, suggesting that most transcriptional changes are not inherited (Figs 7C–D). We found no significant interaction effect between the rearing and ancestral habitats, indicating that the effects of these factors are also independent and additive (Fig. 7E).Pairwise comparisons further supported these findings with the most differentially expressed genes (DEGs) found between scaffold:scaffold and agar:agar groups, and the least between worms of matching growing habitats (Figs 7F–K). Overall, most DEGs genes were uncharacterized or poorly studied to date.

To understand the functional implications of these transcriptional changes, we performed an over-representation analysis (ORA) using gene ontology (GO) slim terms, providing a high-level overview of the affected biological functions. Only 7 GO slims were significantly identified across all groups (Fig. 8). All were upregulated in scaffold-grown animals except for “organelle” and “cell differentiation” which tended to be downregulated (Fig. 8). A complete list of ORA on granular GO terms is also available (Figs S5–7).

**Figure 8.**
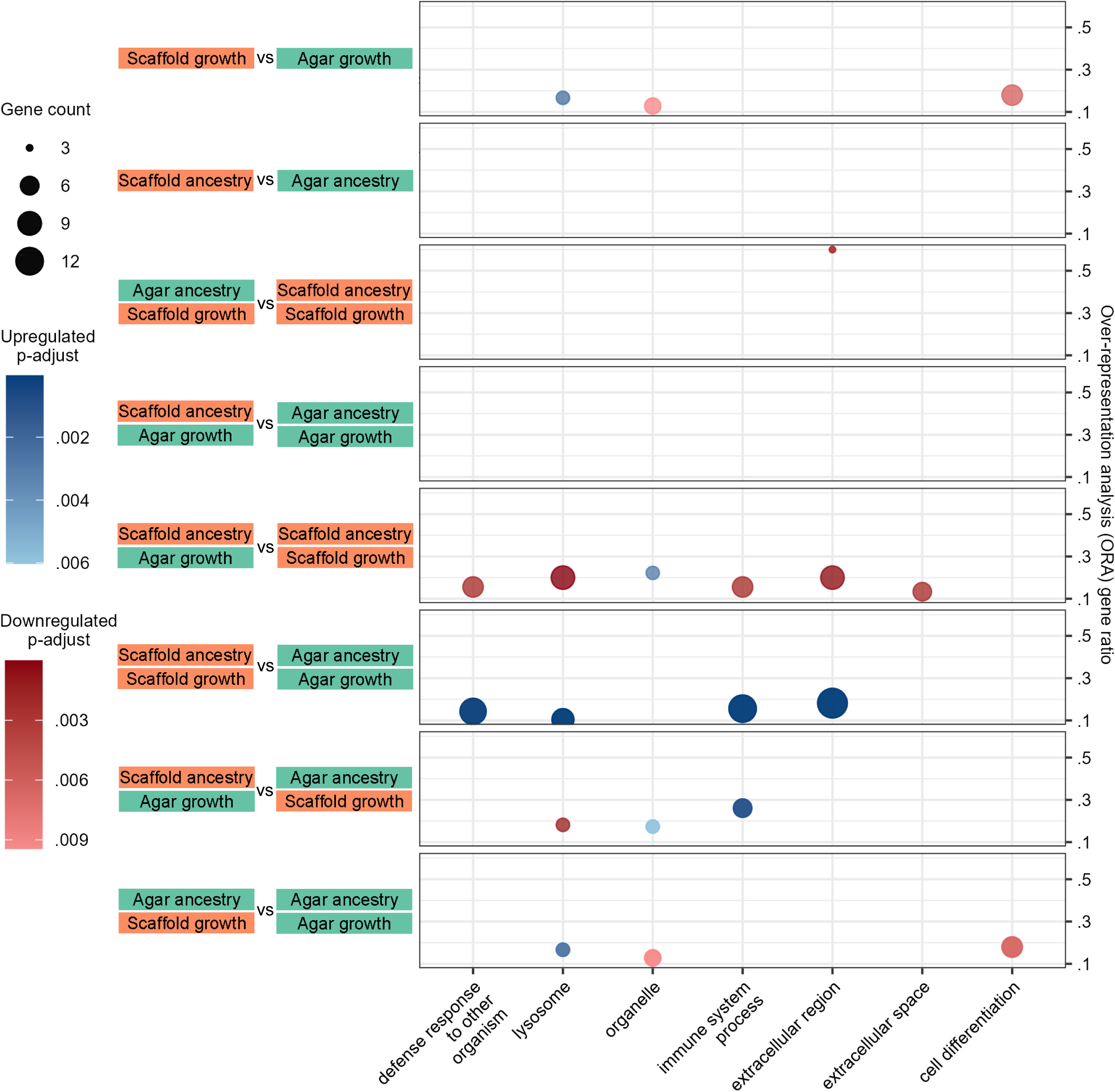
Functional analysis highlights upregulation and downregulation in scaffold-grown worms. Dot plot of over-representation analysis (ORA) of gene ontology (GO) slim terms that were found to be significantly up- or downregulated in the identified differentially expressed genes (DEGs) (Fig. 7C–K) per condition. The following conditions did not have enough DEGs for ORA: ancestry factor (up- and downregulated genes), agar:scaffold versus scaffold:scaffold (upregulated genes), scaffold:scaffold versus agar:agar (downregulated genes), scaffold:agar versus agar:agar (up- and downregulated genes).

## Discussion

This exploratory study sought to investigate the influence of environmental context on various phenotypic and behavioral traits of *Caenorhabditis elegans*. By comparing worms cultured in three-dimensional decellularized fruit tissue scaffolds with those raised under standard two-dimensional agar plate conditions, we aimed to uncover how environmental context affects organismal biology and to set the groundwork for future research in this area. Our preliminary findings suggest that some biological traits of *C. elegans* are affected by the rearing habitat, supporting the notion that laboratory conditions can influence experimental outcomes and potentially our understanding of the studied phenomena.

### Life history modulations

We observed that scaffold-grown worms, regardless of ancestry, had generally reduced body sizes, held a greater number of eggs in their gonads relative to their body size in adulthood, and laid more elongated eggs. Only scaffold-grown worms with also a scaffold ancestry had reduced total brood size. Fat content through life and adult pumping rate were not affected. In previous work, *C. elegans* has been observed to alter its growth rate and brood size with temperature, food availability, oxygen content, population density and presence of pathogens (Byerly et al., 1976; Cypser et al., 2013; Fenk & de Bono, 2017; Harvey & Orbidans, 2011; Harvey & Viney, 2007; Lenaerts et al., 2008; Szewczyk et al., 2006; Wong et al., 2020), but to much greater extremes than what has been observed here. It is crucial to emphasize that our observed life history modulations do not appear to be driven by caloric restriction (CR) as they contrast sharply with that of worms undergoing CR and our RNA sequencing data did not reveal characteristic transcriptional profiles of CR (Fig. S8) (Bar et al., 2016; Lenaerts et al., 2008; Szewczyk et al., 2006; Zhang & Mair, 2017). Overall, our results align with the concept of developmental plasticity, where organisms adjust their life history traits in response to environmental cues (West-Eberhard, 2003). For example, it has been suggested that reduced fecundity in *C. elegans* is beneficial to the colony’s overall fitness (Galimov & Gems, 2020). Our preliminary observations suggest that *C. elegans* can exhibit hierarchical plasticity in its life history in response to environmental changes without the presence of explicit stressors: some traits, such as body size, show greater flexibility and are more readily adjusted, while others, like fat content and reproductive output, demonstrate higher stability, suggesting they may be more critical to the organism’s fitness and are thus more robustly maintained across different environments.

### Behavioral changes in response to environmental context

While most measured behavioral changes were unaffected, we recorded for scaffold-grown animals mild increases in average swimming speed, exploration distance, oxidative stress resistance, and burrowing ability. Our observed behavioral changes could reflect enhancement in neuromuscular abilities or altered sensory processing. These results are similar to research in other model organisms showing that experience with a more complex environments can enhance motor skills, resistance to stressors and disease, and cognitive abilities (Chen et al., 2018; Hillenmeyer et al., 2008; Laviola et al., 2008; Mallory et al., 2016; Nithianantharajah & Hannan, 2006; van Praag et al., 2000; Volgin et al., 2018). While more research is needed to determine how experience with the scaffold habitat affects worm health, our preliminary results suggest that the rearing environment is sufficient to affect some aspects of fitness.

### Transcriptomic profiles of environmental adaptation

Consistent with the relatively subtle phenotypic changes observed, RNA sequencing revealed that the rearing environment induced only mild changes in the transcriptional profiles of young adult worms. The transcriptional adaptations to the environment were highly plastic, arising and reversing within a single generation. Functionally, the DEGs in scaffold-grown animals showed an upregulation in pathways related to defense responses and immune system processes, and a downregulation in genes associated with cell differentiation and organelles. This analysis provides a starting point for understanding the molecular mechanisms underlying environmental adaptation and suggests that while *C. elegans* rapidly adjusts its transcriptome in response to immediate environmental context, many of the affected genes remain poorly characterized.

### Intergenerational inheritance of environmental context

Some measurements differed more drastically in worms that were raised in conditions mismatched from their ancestors, suggesting a possible intergenerationally inherited influence. Particularly, both agar:scaffold and scaffold:agar worms were more quiescent when swimming and exhibited more tortuous crawling paths in the absence of food. This indicates that the environmental mismatch may trigger specific adaptive behaviors under resource scarcity which promote local search. Also, agar:scaffold worms showed reduced success in the burrowing assay, and did not have the reduced brood size observed in scaffold:scaffold worms. Yet, this group did have a generally reduced body size and a similar proportion of eggs in their gonads, suggesting that they could be laying eggs faster. These observations are consistent with studies showing that *C. elegans* can transmit information across generations (Rechavi & Lev 2017). Interestingly, two of the only four DEGs by ancestry (Y102A5C.5 and Y102A5C.6) are pseudogenes that have been found to be involved in transgenerational inheritance and learning (Posner et al., 2019; van der Linden et al., 2010). This strongly suggests that the observed behavioral inheritance might be mediated specifically through small RNAs and post-translational modifications which are not detected by traditional mRNA sequencing. Nonetheless, the rapid reversal of the vast majority of gene expression changes and phenotypes indicates that most of the observed effects are primarily due to direct environmental influences rather than intergenerationally inherited modifications. This capacity for rapid adaptation and equally rapid reversal is consistent with the natural boom-and-bust, transient lifestyle of *C. elegans* (Frézal & Félix 2015).

The differences we have observed suggest that standard laboratory conditions may not fully capture the range of natural phenotypic and genetic variability in *C. elegans*. By demonstrating that *C. elegans* can modify their reproductive strategies, development, and behavior in response to habitat context, we also underscore this organism’s capacity for phenotypic plasticity, which is a key factor in evolution and adaptation. Our results highlight the sensitivity of biological research to environmental context, reminding us of the importance of proper controls and reproducibility in scientific research.

As an exploratory study, the sample sizes for certain assays may not be sufficient to draw definitive conclusions; larger cohorts would increase statistical power and confidence in the results. While the scaffold habitat provides structural enrichment, other environmental factors, such as differences in local food availability, oxygen diffusion, or waste accumulation, could be contributing to the observed effects. Controlling for these factors or understanding their dynamics in the scaffold would strengthen the causal links between environmental complexity and phenotypic outcomes. For example, variations in the occupancy task (Guisnet et al., 2021a) could help elucidate which characteristics make the scaffold habitat attractive. While we identified DEGs, the specific molecular pathways mediating the observed phenotypic changes require further investigation through targeted genetic manipulations, epigenetic modifications, and the study of inherited small RNAs and proteomic profiles. Likewise, a large inventory of additional physiological parameters still remains to be evaluated.

In conclusion, this exploratory study demonstrates the impact of environmental context on *C. elegans* biology, challenging the notion that findings from simplified laboratory conditions can fully capture the organism’s biological potential. By bridging the gap between laboratory and more naturalistic conditions, this work sets the stage for a more nuanced understanding of gene-environment interactions in *C. elegans* and potentially other model organisms. Future research building on these findings could lead to significant advances in our understanding of phenotypic plasticity, adaptation, and the interplay between genes and environment in shaping biological outcomes.

## Materials and Methods

### Strains and culture

Wild-type N2 *C. elegans* provided by the CGC were grown under standard conditions at 20 C° and were fed OP50 *E. coli* bacteria (Brenner, 1974).

Scaffold and traditional agar plate habitats were created as described in Guisnet et al. (2021b). Animals were first thawed from fresh wild-type N2 stock and maintained for 10 generations on traditional agar plates. Then, maintenance on scaffold plates started in parallel for at least 10 generations before experimental testing.

### High-resolution magic angle spinning (HR-MAS) nuclear magnetic resonance spectroscopy

The scaffolds were prepared as described in Guisnet et al. (2021b) up until storage at 4℃. 30 samples were taken out of 5 different stored scaffold slices using a biopsy punch. The samples were left to vacuum dry for 5 hours at room temperature and then rehydrated in deuterated water (D_2_O, D 99.9%).

HR-MAS NMR experiments were performed on a Bruker Avance III HD spectrometer operating at a resonance frequency of 599.90 MHz for 1H. The instrument was equipped with a 4 mm HR-MAS dual inverse 1H/13C probe with a magic angle gradient. All experiments were carried out at a magic angle spinning rate of 5 kHz and a temperature of 298 K.

Bruker TOPSPIN software (version 3.6, patch level 5) was used to acquire and process the NMR data. The 1D 1 H HR-MAS NMR spectra were recorded using a 1D NOESY two-step presaturation sequence for water suppression (“noesypr1d” from the Bruker pulse-program library). Each 1D 1 H NMR spectrum consisted of co-adding 256 transients with a spectral width of 12 kHz, a data size of 12K points, an acquisition time of 0.5 second, and a relaxation delay of 2 second. The coadded free induction decays (FIDs) were exponentially weighted with a line broadening factor of 0.3 Hz, Fourier-transformed, phase and (polynomial) baseline corrected to obtain the 1 H NMR spectra.

### Pumping rate

Pumps of young adults were counted directly on maintenance plates at room temperature on a compound microscope. 10 minutes before counting, 3 2 µL drops of IAA diluted 1:100 in double-distilled water were added to the lid of the plates to encourage movement to the surface. The experimenter counting the pumps was blind to the ancestry.

### Developmental timing, gonad egg count and egg size

Worms were timed by placing egg-laying adults on plates for 2 hours. These offspring were moved to fresh plates through the experimental days as needed to keep them fed. Every 24 hours after being laid, worms were placed on 2% agarose pads and immobilized with 1M sodium azide and imaged within 10 minutes. Grayscale images were taken with an Olympus XM10 CCD camera mounted on an Olympus MVX10 microscope using an MV PLAPO 2XC objective. TIF images were acquired with a 1376 × 1038 resolution and 10 ms exposure with the cellSens software (Evident).

Starting on day 3, two images were taken for each animal: one with the head and tail in focus, and a second with the mid-body in focus. On day 5, adults were moved to an empty agar plate to lay eggs which were used for imaging

Worm and egg images were segmented as described in Guisnet & Hendricks (2025). Egg elongation was determined by fitting an ellipse to the egg contour and calculating the ratio of the major axis to the minor axis.

The number of eggs present in the worm gonads were counted with the images that were focused on the midbody of the worm by a *C. elegans* expert blind to the condition using the LabelBox web tool with academic access (*Labelbox*, n.d.).

### Oil Red O fat staining

Oil Red O staining was done following the protocol described in He (2012) except that instead of using a thermomixer shaker, tubes were taped to a box filled with room-temperature heat beads and shaken at 150 RPM.. For each group, worms were pooled from differently aged well-fed plates to obtain a mix of stages. Colored images were taken with a QImaging MicroPublisher RTV 3.3 camera with a Leica 10445930 1.0x mount lens. TIF images were acquired with a 2048 x 1536 resolution, 75 ms exposure and 24-bit depth. Worms were segmented as described in Guisnet & Hendricks (2025).

To quantify stain intensity, the RGB channels of worm images were first normalized using background pixel values. A composite staining metric was calculated for each pixel by multiplying the red dominance, which measures the intensity of red relative to green and blue, by the inverse luminance to account for the darkening effect of the stain. Overall worm staining was calculated as the sum of the staining metric for each pixel divided by the worm area in pixels.

### Brood size

The same protocol as described in Guisnet et al. (2021b) was followed to create both habitats, with the following modifications: 4 cm diameter plates were used, scaffold were cut to 1.5 cm diameter, 10 µL of NGML *E. coli* culture was added directly on both types of plates, plates were parafilmed and kept upside down in a closed box 24 hours prior to use.

Worms were synchronized and placed on individual plates at the late L4 stage. Every 24 hours, for 4 days, worms were moved to a fresh plate to distinguish egg-laying days. Worms that died or found to be in plates with contamination were completely removed from the dataset. Offspring were counted 3 days after being laid. Scaffold-born offspring were lured out of the scaffold with food, and manually extracted from the scaffold by shaking in liquid.

### Burrowing assay

We performed the burrowing assay as described in Laranjeiro et al. (2019) with minor modifications. We used 96-well plates and added only one worm per well. 5 µL of LB *E. coli* OP50 culture was used as the attractant at the top of the pluronic gel. Young adult worms were selected from maintenance plates and left to roam off food for 2 minutes before being added to the well.

### Oxidative stress assay

Young adult worms were selected from maintenance plates and moved to empty agar plates to crawl off food for 5 to 10 mins. 180 µL of 3 mM H_2_O_2_ was added to each well of a 96-well plate and 1 worm per well was transferred with a platinum worm pick. Worms were checked every 2 hours for 8 hours. Before marking an inactive worm as dead, it was gently poked with a platinum worm pick. Animals were also checked at time 0 (right after being added to the well). Animals dead at time zero were removed from the dataset.

### Long-term swimming bends count

This protocol was inspired by (Ghosh & Emmons, 2008). Young adult worms were picked from maintenance plates and left to roam off food for 10 minutes on an empty agar plate. They were then transferred to individual wells of a 96-well plate filled with 180 µL of room temperature filtered NGML. Counting started within 10 minutes of the first worm being transferred. The number of bends made in 5 seconds was counted by eye with an auditory timer every 5 minutes over 3 hours. Worms that were found dead at the end of the experimental period were removed from the dataset.

For high viscosity swimming, 0.5% and 1% methyl cellulose (MC) in filtered NGML was prepared by serial dilution. Their viscosity was verified with a RheoSense m-VROC II viscometer to be 5.6 cP (similar to milk) and 20 cP (similar to vegetable oil), respectively. The viscosity of filtered NGML was verified to be 1 cP (water).

### Short-term droplet swimming recording and analysis

50 µL of filtered NGML or 3.5 µL of MC solution was pipetted onto the center of a 10 cm diameter plexiglass dish. Young adult worms were selected from maintenance plates and left to roam off food on an empty agar plate for 2 minutes before being moved to the droplet with a platinum worm pick.

AVI videos were recorded for 1 minute in NGML and 30 seconds in MC solutions at 10 fps with a FLIR Blackfly (BFS-U3-51S5M-C) 5.0 MP camera and Spinnaker SDK software (Teledyne) at 2448 x 2048 resolution. The recordings were analyzed as described in Guisnet & Hendricks (2025).

### Crawling recording and analysis

Young adult worms were selected from maintenance plates and left to roam off food on an empty agar plate for 2 minutes. 10 cm diameter NGM agar plates were used with either nothing (no food), 300 µL of NGML OP50 *E. coli* culture added 1 hour before recording (low food density) or 300 µL of NGML OP50 *E. coli* culture incubated at 37℃ for 24 hours and acclimated at room temperature prior to recording (high food density).

AVI videos were recorded for 1 minute as described for swimming and analyzed as described in Guisnet & Hendricks (2025).

### RNA sequencing and data processing

Young adult *C. elegans* were collected by hand from maintenance plates directly into 25 μL of TRIzol reagent. Samples were stored at -80°C until sufficient quantities were collected.

Frozen samples were vortexed for 15 minutes and 150-250 worms were pooled for each sequencing sample. These pooled samples underwent three freeze-thaw cycles: freezing at -80°C for at least 20 minutes, followed by 15 minutes of vortexing before final storage at -80°C until sequencing.

Samples were further processed and sequenced by the GenomeQuébec genomics facility. Total RNA was extracted and sequenced using Illumina NovaSeq 6000 platform, generating paired-end reads of 100 bp. Three replicates per condition were sequenced, with each replicate receiving 2 x 25 million reads.

Sequencing reads were cleaned and aligned to the *C. elegans* reference genome using a custom Linux shell script. Quality control of raw sequencing data was performed using FastQC (v0.11.9) and MultiQC (v1.12) (Andrews, 2010; Ewels et al., 2016). Reads were trimmed using Trimmomatic (v0.39) to remove low-quality bases and potential adapter contamination (Bolger et al., 2014). Specifically, the first 10 bases were removed from the 5’ end of each read to mitigate potential priming biases, and a sliding window quality trimming was performed (window size 4, required quality 15). Trimmed reads shorter than 36 bp were discarded. The trimmed high-quality reads were then aligned to the *C. elegans* reference genome (WBcel235) using HISAT2 (v2.2.1) with default parameters (Kim et al., 2019). Post-alignment, a minimal filtering step was performed to remove a small number of malformed alignment entries (31 out of 161,930,474 total entries). The resulting alignments were converted to BAM format, sorted, and indexed using SAMtools (v1.13) (Danecek et al., 02 2021). Gene-level quantification was performed using featureCounts (v2.0.3) against the Ensembl gene annotation (release 104) (Liao et al., 2014). Further analysis was performed in R (v4.3.2) (R Core Team, 2021).

Gene names, gene ontology (GO) terms, GO slim terms and all their corresponding descriptions were fetched from WormBase BioMart with the biomaRt package (v2.58.2) using the “celegans_gene_ensembl” dataset (Durinck et al., 2009; Harris et al., 2020; *WormAtlas Navbar*, n.d.). Genes were matched to GO terms with the org.Ce.eg.db package (v3.18.0) (Carlson, 2023).

Differential expression analysis was performed with the count data using the DESeq2 package (v1.42.1) (Love et al., 2014). A DESeqDataSet object was created using the count data and condition labels (pairwise and factor-wise), and differential expression analysis was conducted using the DESeq function with default parameters. PCA data was generated with the DESeq2 plotPCA function. Volcano plots were generated with the EnhancedVolcano package (v1.20.0) (Blighe et al., 2023). Over-representation analysis was conducted on GO and GO slim terms with enrichGO and enricher function of the ClusterProfilter package (v4.10.1) with default parameters (Wu et al., 2021).

### Statistical analysis

Image analysis was conducted in Python (v3.12.3), and all data manipulation and visualization were conducted in R (v4.3.2) with the tidyverse (v2.0.0) and ggplot2 (v3.5.2) (R Core Team, 2021; *The Python Language Reference*, n.d.; Wickham, 2016; Wickham et al., 2019). All boxplots represent the interquartile range (IQR), with the median indicated by the horizontal line. Whiskers extend to the furthest data point within 1.5 times the IQR. Outliers beyond this range are shown as individual points. Scatterplots have all data points shown.

Given the exploratory nature of this study, we did not perform formal hypothesis testing or compute p-values. Instead, we focused on estimating effect sizes and calculating confidence intervals to assess the magnitude and precision of observed associations. This approach aligns with our objective to generate hypotheses and uncover novel insights without the constraints of predefined hypotheses.

By not computing p-values, we aimed to minimize the risk of Type I errors that can arise from multiple comparisons in exploratory analyses. This strategy reduces the likelihood of identifying spurious effects that may occur by chance when testing numerous variables. Furthermore, avoiding p-value calculations helps prevent the misinterpretation of statistical significance in a context where findings are preliminary and require validation in future confirmatory studies (Makin & Orban de Xivry, 2019).

## Acknowledgements

We thank Stephanie C. Weber from McGill University for sharing their colored camera for Oil Red O imaging. Christopher Moraes and Claire Edrington for the viscosity measurements. Giovanni Vizcardo for helpful insights on worms in viscous liquids. WormBase and WormAtlas teams for providing invaluable resources. All members of the lab for helpful discussions and comments. Strains were provided by the *Caenorhabditis* Genetics Center (CGC), which is funded by NIH Office of Research Infrastructure Programs (P40 OD010440).

## Data and resource availability

RNA sequencing data will be available on NCBI GEO and all other datasets are available on FigShare (10.6084/m9.figshare.30071044).

