## Supplementary figures and images for "The impact of rearing environment on *C. elegans*: Phenotypic, transcriptomic and intergenerational responses to 3D enriched habitats"

### Supplemental Figure 1

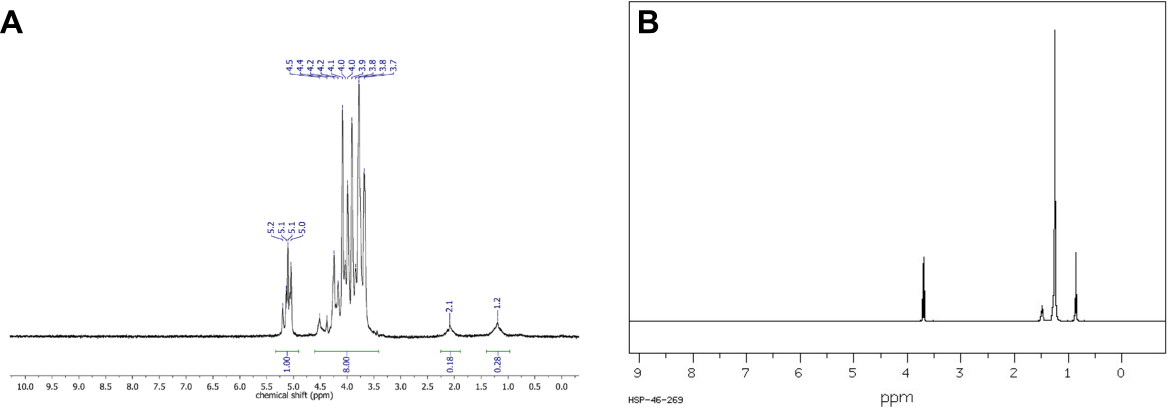

### Supplemental Figure 2

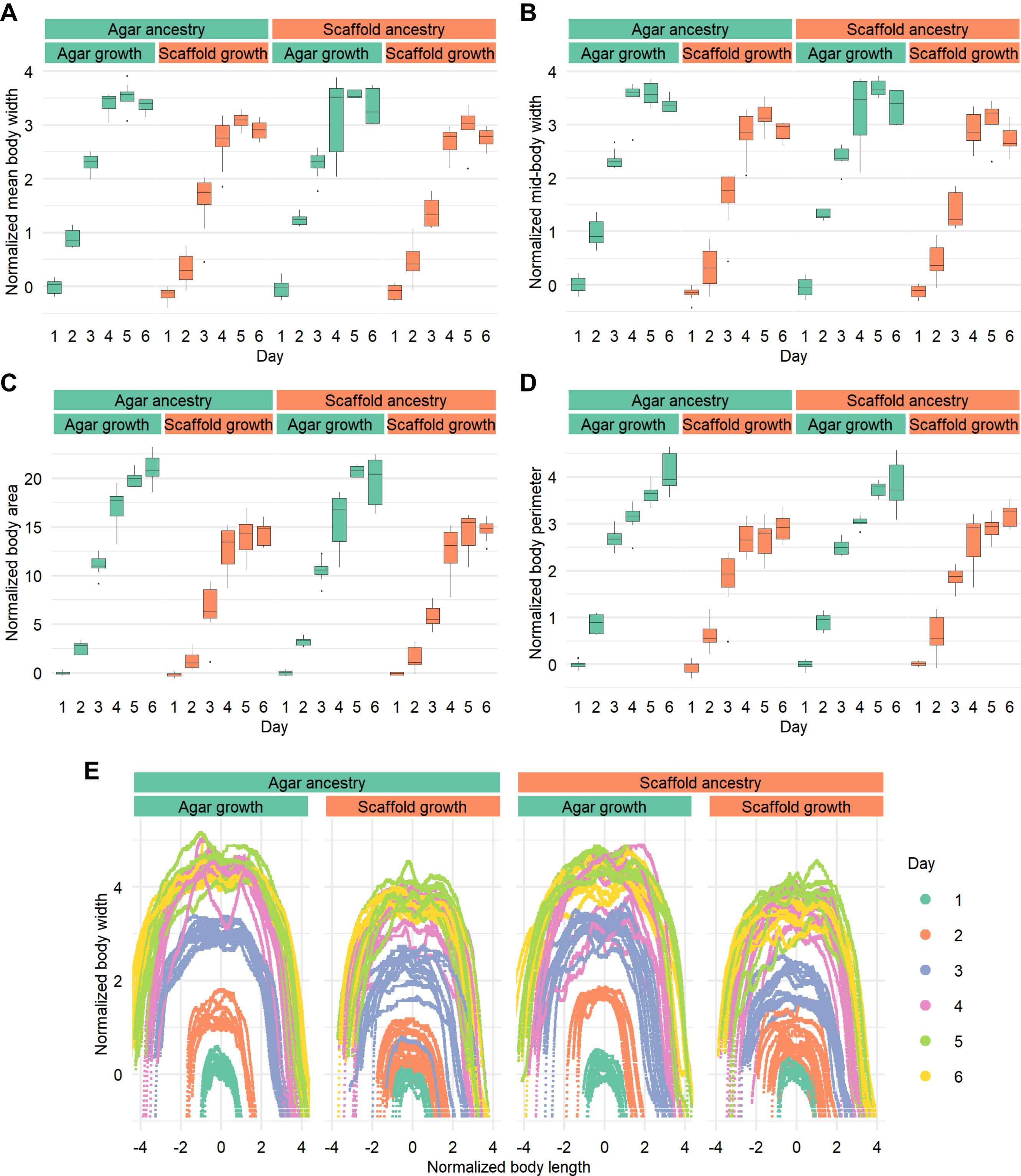

### Supplemental Figure 3

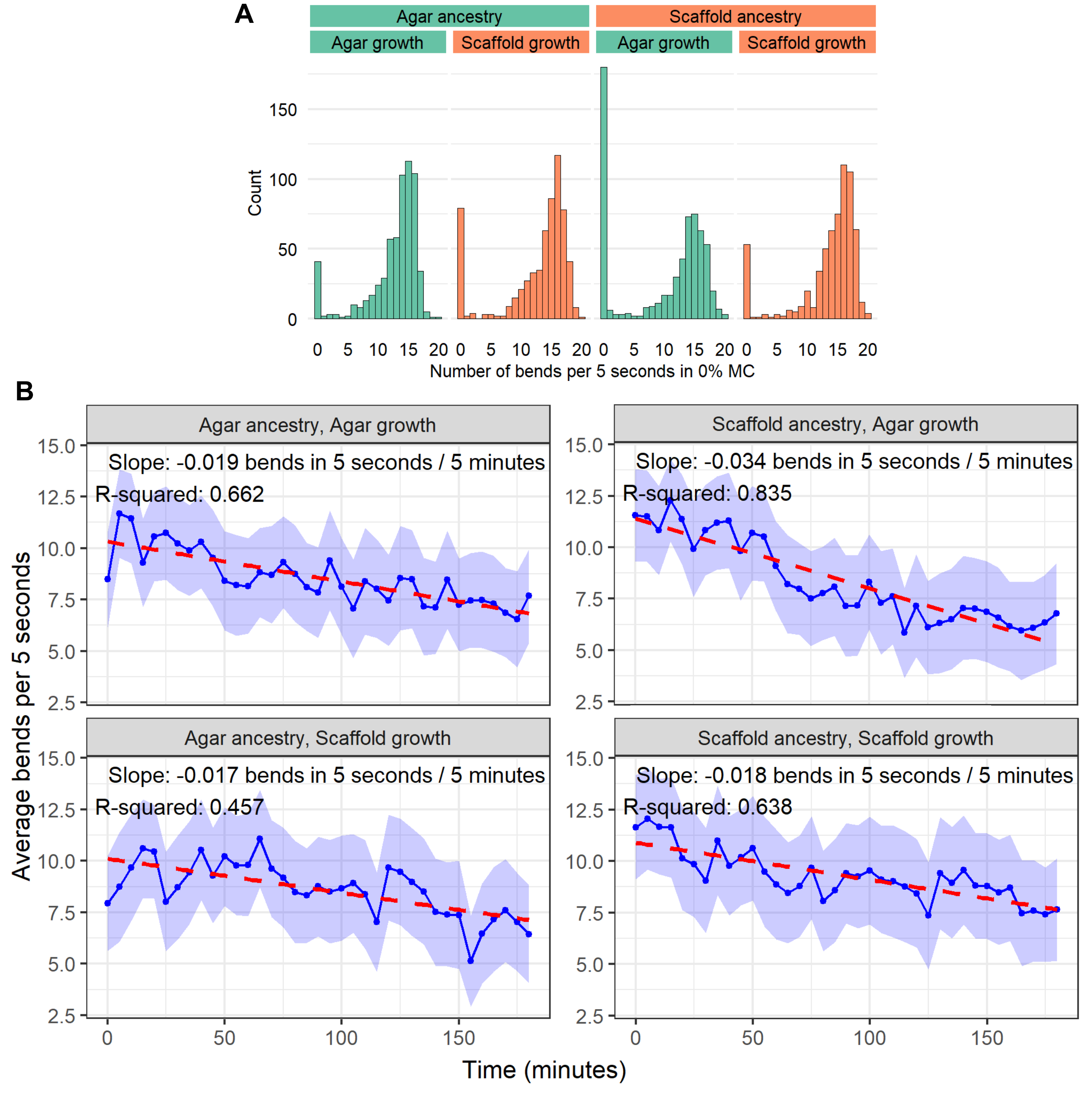

### Supplemental Figure 4

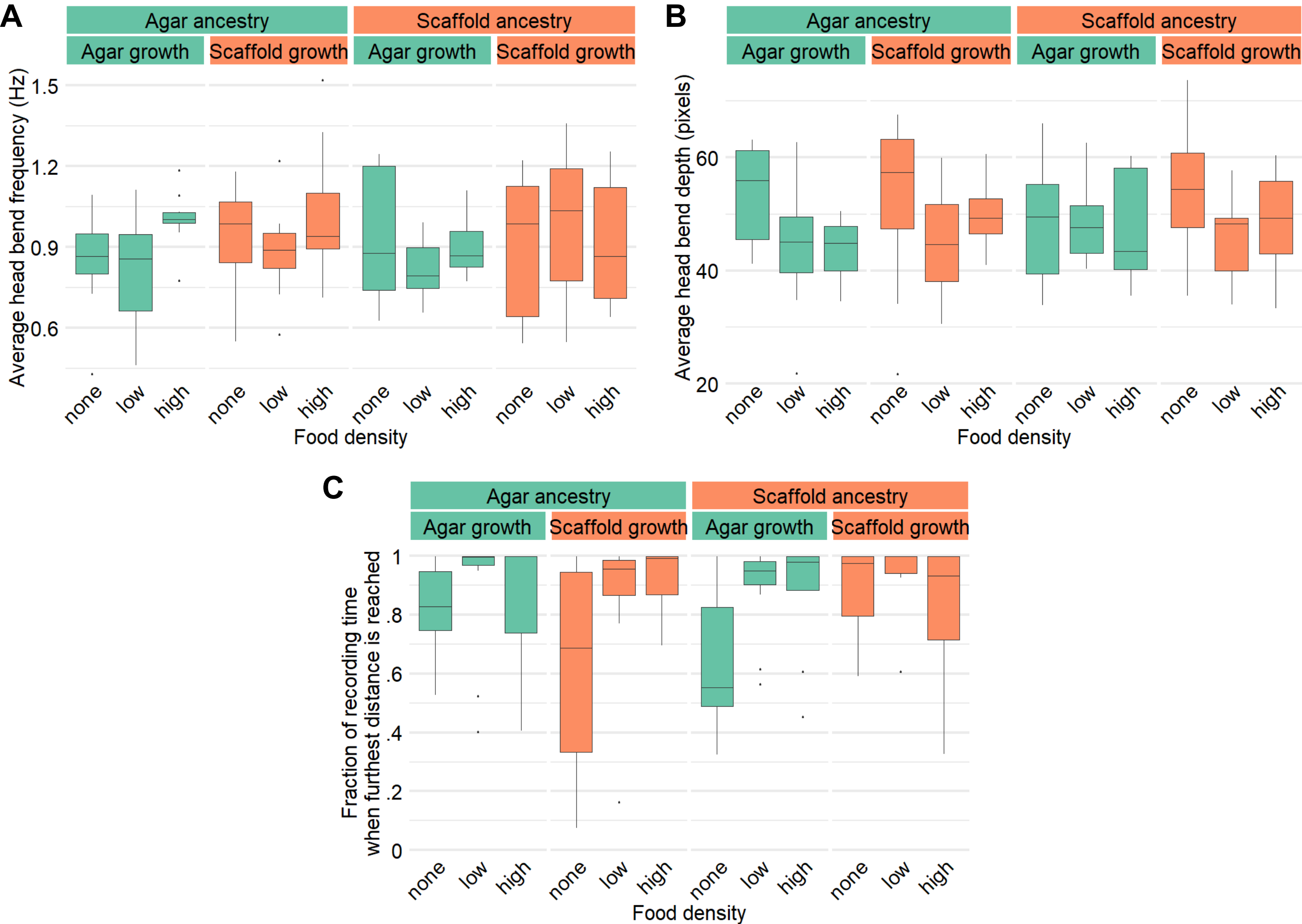

### Supplemental Figure 5

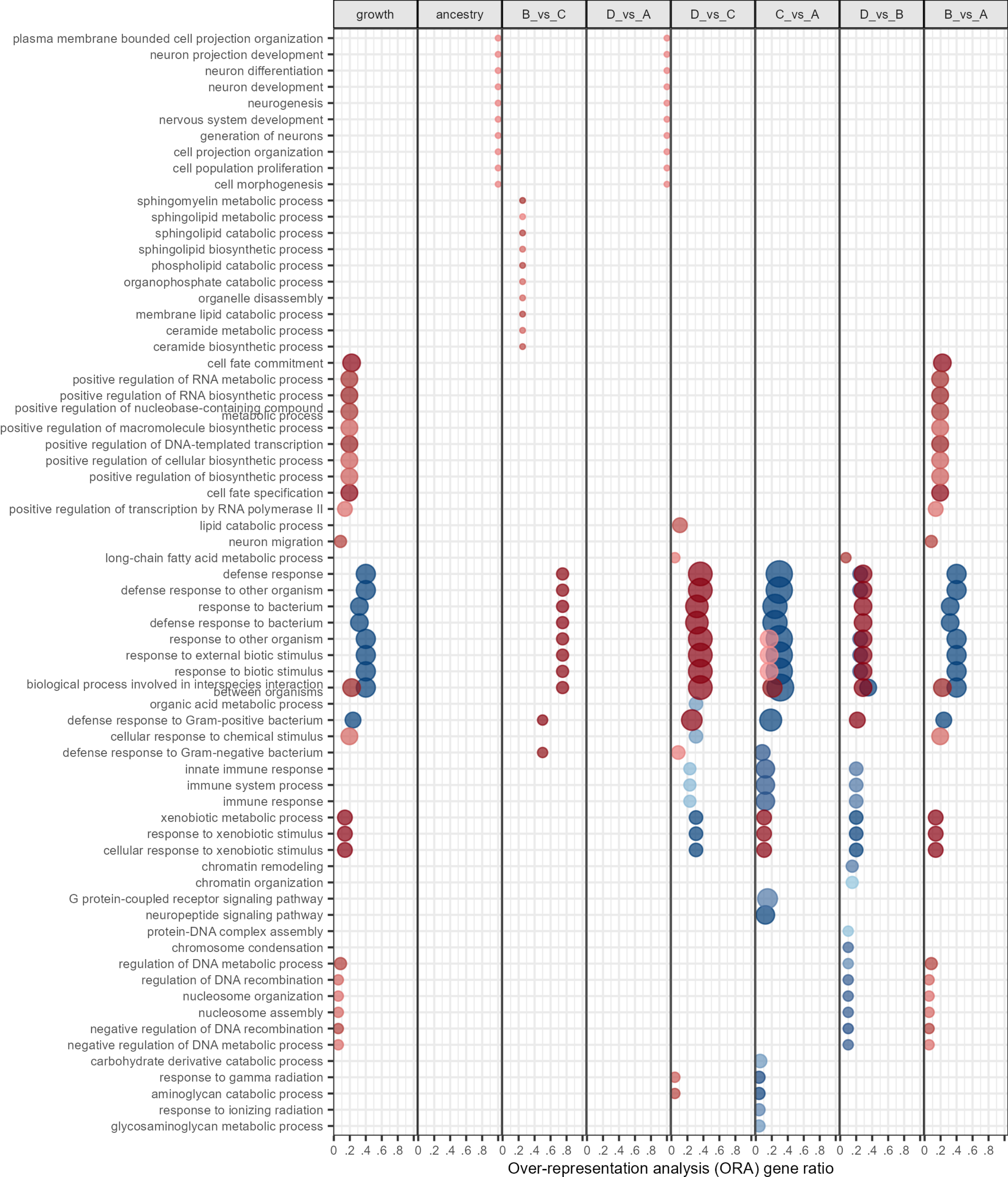

### Supplemental Figure 6

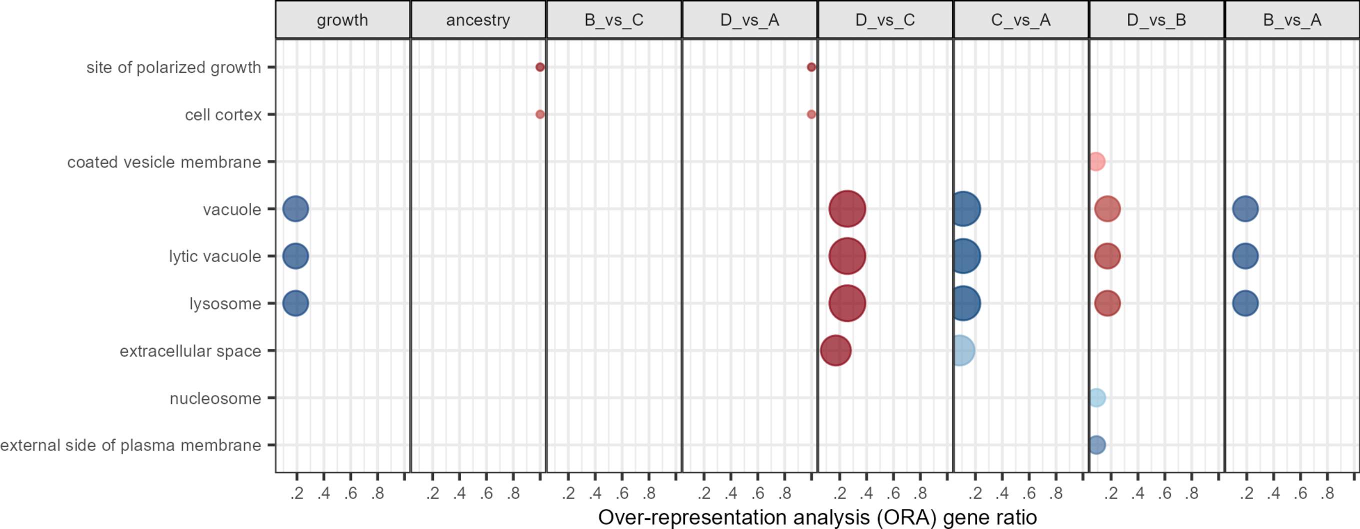

### Supplemental Figure 7

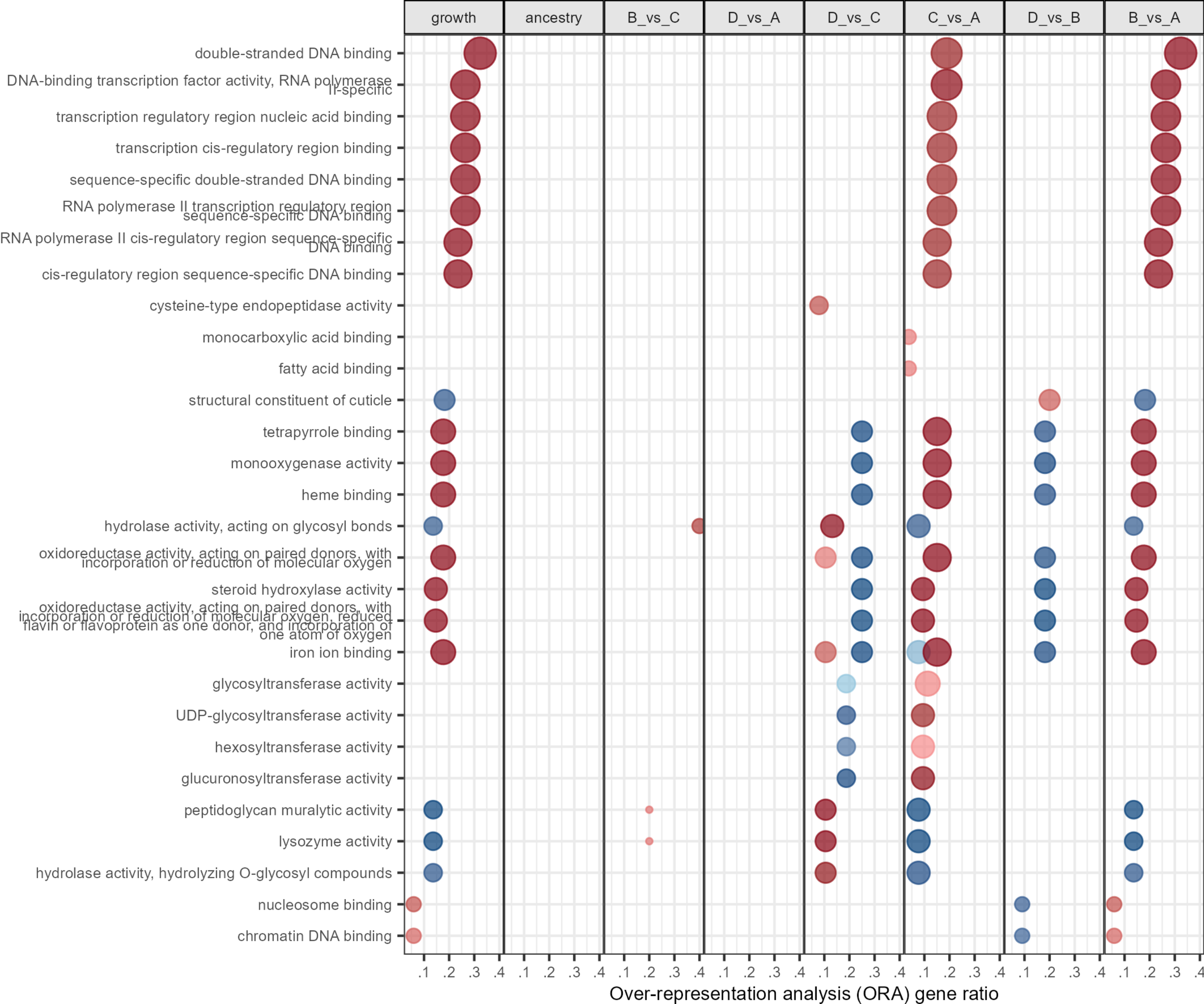

### Supplemental Figure 8

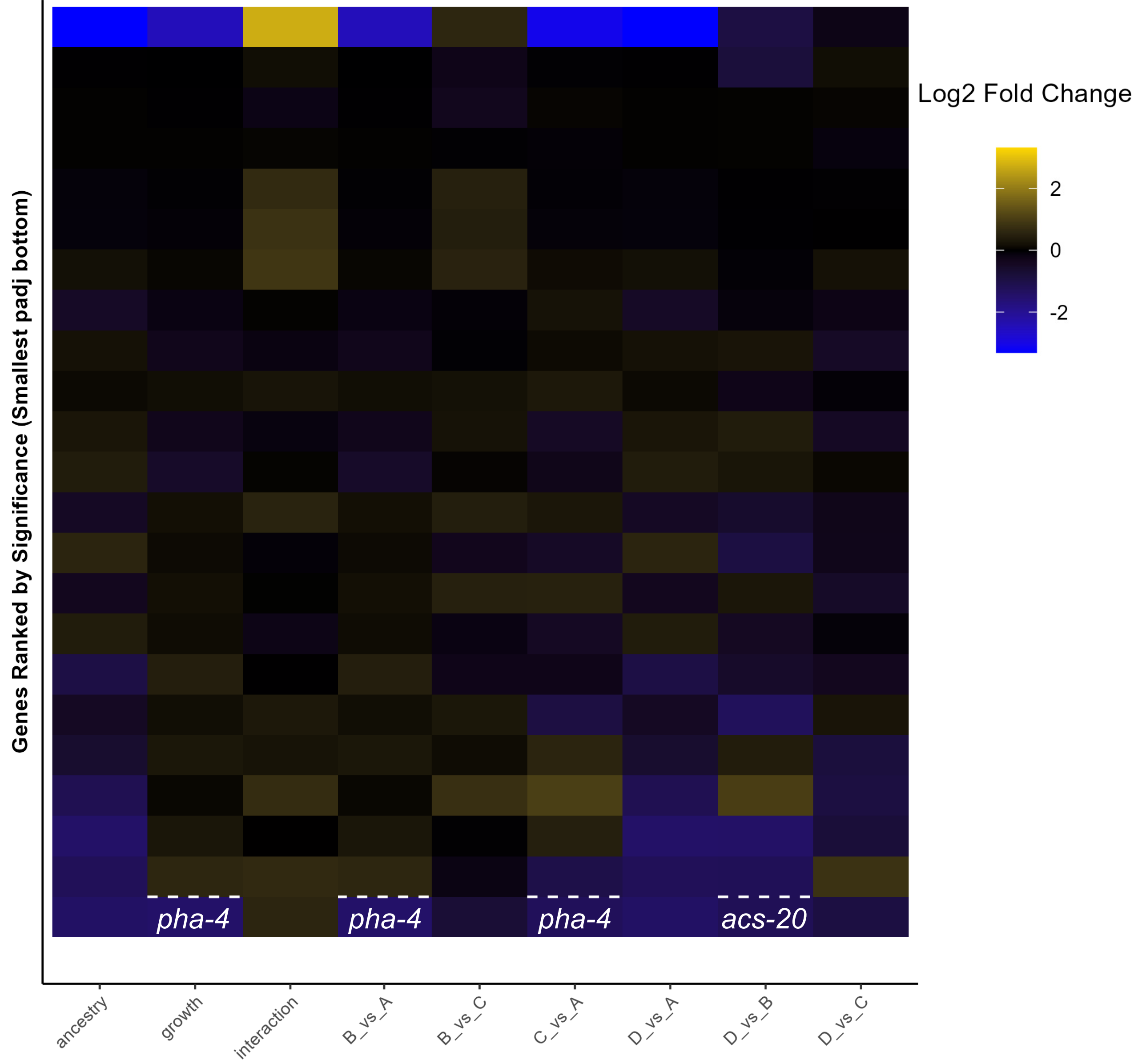
